# In Situ Volumetric Imaging and Analysis of FRESH 3D Bioprinted Constructs Using Optical Coherence Tomography

**DOI:** 10.1101/2021.06.30.450389

**Authors:** Joshua W. Tashman, Daniel J. Shiwarski, Alexander Ruesch, Frederick Lanni, Jana M. Kainerstorfer, Adam W. Feinberg

## Abstract

As 3D bioprinting has grown as a fabrication technology, so too has the need for improved analytical methods to characterize these engineered constructs. This is especially challenging for soft tissues composed of hydrogels and cells as these materials readily deform, posing a barrier when trying to assess print fidelity and other properties non-destructively. Indeed, given the importance of structure-function relationships in most tissue types, establishing that the 3D architecture of the bioprinted construct matches its intended anatomic design is critical. Here we report development of a multimaterial bioprinting platform with integrated optical coherence tomography (OCT) for in situ volumetric imaging, error detection, and 3D reconstruction. While generally applicable to extrusion-based 3D bioprinting, we also report improvements to the Freeform Reversible Embedding of Suspended Hydrogels (FRESH) bioprinting process through new collagen bioink compositions, support bath optical clearing, and machine pathing. This enables high-fidelity 3D volumetric imaging with micron scale resolution over centimeter length scales, the ability to detect a range of print defect types within a 3D volume, and real-time imaging of the printing process at each print layer. These advances provide FRESH and other extrusion-based 3D bioprinting approaches with a comprehensive methodology for quality assessment that has been absent in the field to date, paving the way for translation of these engineered tissues to the clinic and ultimately to achieving regulatory approval.

**Teaser:** Transparent FRESH support bath enables in situ 3D volumetric imaging and validation of patient-derived tissue constructs.

## Introduction

As the demand for organ transplantation continues to outpace the supply, clinicians and researchers are turning to regenerative medicine and tissue engineering strategies to create tissue *de novo* (*1*). 3D bioprinting has emerged as a way to build these tissues using robotic control to precisely pattern cells and biological hydrogels in a layer-by-layer process (*2*). This technology has been used to produce a number of advanced constructs including heart valves, cardiac tissue scaffolds, perfusable vascular networks, kidney proximal tubules, external ear scaffolds, and model lung alveoli (*3–9*) for applications ranging from in vitro models to tissue replacements (*4, 5, 10–12*). As bioprinting capabilities improve and constructs become larger and more geometrically complex, there is an increasing need for a hardware and software platform enabling volumetric 3D imaging and validation. Existing imaging techniques allow for characterization of construct fidelity after printing; however, by acquiring images during the printing process we can create full 3D reconstructions of geometrically complex prints and monitor the fabrication process in ways that were previously impossible.

While an integrated platform for bioprinting and volumetric imaging is valuable for all bioprinting modalities, it is even more useful for embedded bioprinting techniques such as Freeform Reversible Embedding of Suspended Hydrogels (FRESH). In FRESH, the support bath is critical to enabling true freeform printing, but is highly light scattering making live viewing and imaging of the print process challenging (*3, 4*). Commonly used imaging techniques, such as bright field imaging, confocal fluorescence microscopy, and micro–Computed Tomography (µCT) either lack the ability to capture 3D images, are too slow, or cannot image through the scattering support bath. An ideal imaging solution would need to have a large imaging depth and field of view, a fast volumetric acquisition speed, the ability to resolve features from the micron to centimeter scale, and be non-destructive to the sample. An imaging modality that fulfills these requirements is optical coherence tomography (OCT). OCT has been a valuable tool for clinical imaging in ophthalmology of the retina and cornea (*13–15*), creating 3D angiograms (*16, 17*), and even monitoring fabrication and quality of electronic devices (*18, 19*). More recently, the use of OCT has emerged in the 3D printing space to monitor print quality for simple process feedback control, and to image collagen, PEG, and 3D printed hydrogel scaffolds after printing (*20–25*). While these reports demonstrate advantages of using OCT to characterize a printed scaffold, to achieve true 3D reconstruction of printed constructs with complex internal architecture there remains a need to integrate a 3D imaging system, such as OCT, directly into high-performance 3D bioprinters.

Here we address this challenge by developing a custom-built dual extruder 3D bioprinter with an integrated OCT system to perform real-time imaging during FRESH printing. In addition to developing the custom hardware, we enhanced the OCT contrast of our collagen I bioink, developed a method to image prints larger than the maximum imaging depth (>8.3 mm), and created a transparent support bath to address the problem of FRESH gelatin microparticle support bath opacity. Finally, we demonstrate the flexibility of this platform by imaging live during printing, intermittently between print layers, or following print completion. Together these advancements enable us to (i) visualize our constructs while they are still embedded, providing in situ measurements of the as-printed geometry, (ii) assess the quality of large bioprinted constructs such as human-scale tissues and organs, and (iii) generate 3D reconstructions of the bioprinted construct for dimensional measurements, error detection, and to ensure geometric fidelity.

## Results

### Integration of Optical Coherence Tomography into a High-Performance 3D Bioprinter

Current 3D bioprinters lack real-time monitoring and cannot assess the 3D structure of printed constructs. To address this, we designed and built a high-performance 3D bioprinter with an integrated OCT scan head for in-situ volumetric imaging during printing (Fig. 1A and fig. S1). In addition to the OCT scan head, the bioprinter incorporated two Replistruder 5 syringe pump extruders improved from our previously published Replistruder 4 open-source design (*26*). Performance was significantly improved compared to our previously published open-source bioprinters based on desktop-grade thermoplastic extrusion printers (*5, 27*). While these low-cost printers have adequate performance for a wide range of bioprinting applications (*3–5, 28*); the positional accuracy is limited by their belt and pully driven motion system and is on the order of ±100 µm (*29*). Our new design utilizes a gantry configuration of four high-precision stages and achieves positional accuracy of 8 µm over their full 100 mm travel (*30*).

**Figure 1.**
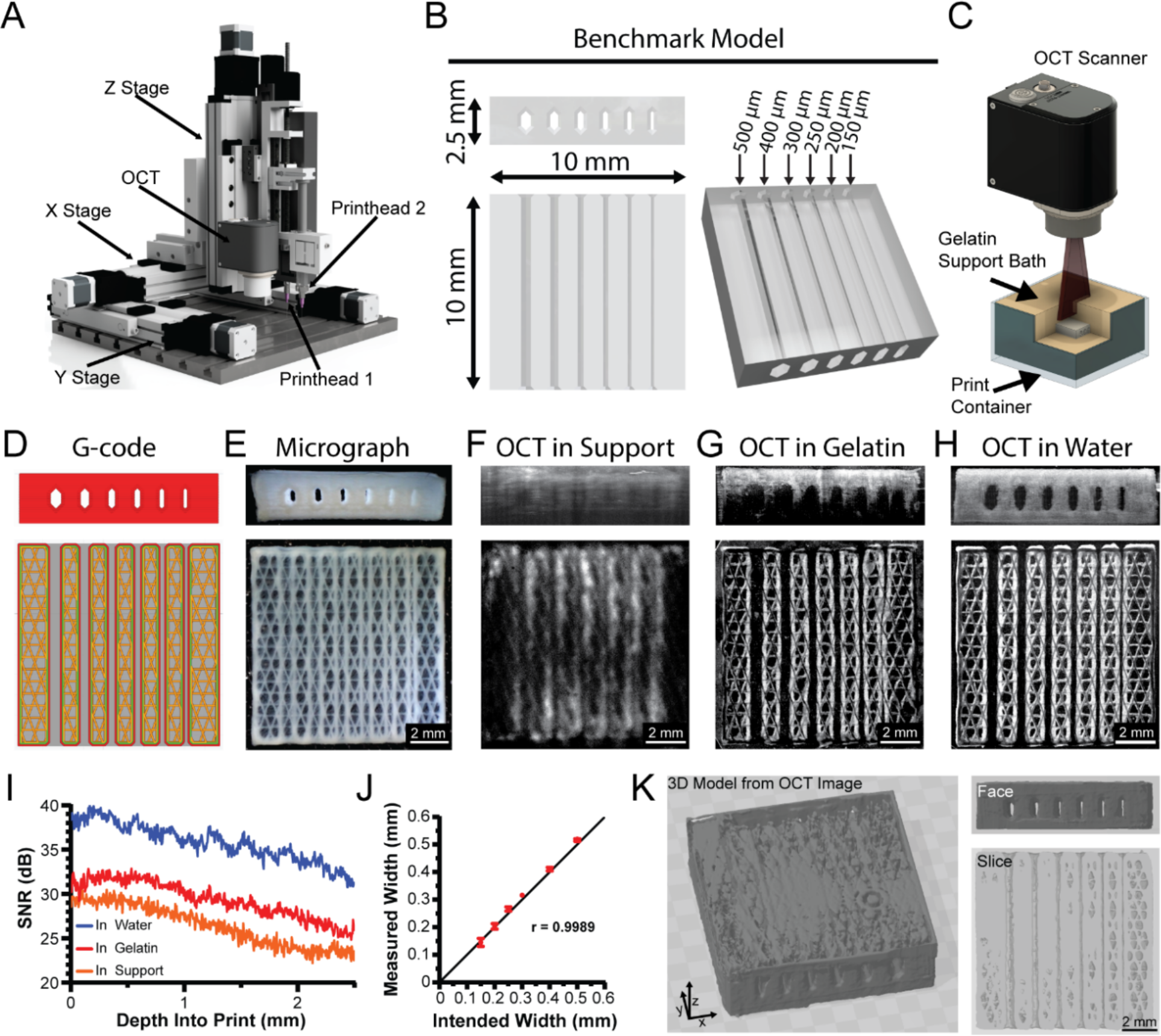
Design and integration of optical coherence tomography (OCT) into the FRESH printing platform. (**A**) 3D render of a custom-designed high-performance 3D bioprinter with dual extruders and an OCT scan head. (**B**) A benchmark model with internal channels of varying widths to test printer performance, resolution capabilities, and imaging quality. (**C**) Example rendering of OCT imaging of a FRESH printed construct within the gelatin microparticle support bath. (**D**) G-code 3D printer pathing of benchmark model channel cross section. (**E**) Photomicrograph of the benchmark model FRESH printed from collagen-I showing end on and top-down views. (**F-H**) OCT imaging of the benchmark model in the gelatin microparticle support bath, melted and resolidified gelatin, or water respectively. (**I**) Analysis of the OCT SNR for the collagen benchmark model within either gelatin microparticle support bath, melted and resolidified gelatin, or water. All three SNR curves are statistically significantly different from each other (p<0.0001, n = 12 total measurements from 3 benchmark prints each, Wilcoxon signed-rank test). (**J**) Correlation between the intended internal channel width and the measured width via OCT (n = 4, Pearson’s correlation of r = 0.9989). (**K**) Images of the 3D OCT image data showing the full reconstruction, an end on view, and a cross-section through the internal channels.

To assess printing resolution and OCT imaging capabilities, we developed a benchmark model with six open channels ranging from 150 up to 500 µm in width, individual filaments as small as 100 µm in diameter, and overall size up to 10 mm (Fig. 1B). This benchmark was designed to be imaged with the integrated OCT scan head during printing while embedded in the FRESH gelatin microparticle support bath (Fig. 1C). For printing, the benchmark model was processed into G-code using open-source slicer software for the bioprinter to execute (Fig. 1D). Once FRESH printed from a collagen type I, the benchmark model was released from the support bath, washed to remove melted gelatin, and imaged with a camera (Fig. 1E). This serves as an example of the standard way 3D printed scaffolds are analyzed using microscopy or photography and while it shows that all channels are visibly open on the exterior surface, it provides minimal information on the internal structure and print fidelity.

Next, we evaluated OCT imaging of the benchmark model (i) in the gelatin microparticle support bath, (ii) in melted and resolidified gelatin, and (iii) in water. We began by imaging the benchmark model while it was still embedded in the highly light-scattering gelatin microparticle support bath (Fig. 1F). The outlines of larger features and the channels are visible, but the fidelity of the image is not sufficient for accurate measurement. To reduce the support bath opacity, we melted the microparticles and resolidified the gelatin before capturing another OCT image (Fig. 1G). The benchmark model’s infill and the channel lumens can now be clearly identified, but the signal decays sufficiently throughout the depth of the print such that the bottom layers cannot be resolved. Finally, we imaged the benchmark model in water (Fig. 1H) and all the channels and features of the G-code are resolved throughout the depth of the print. A side-by-side comparison for the benchmark imaged in the support bath, resolidified gelatin, and water highlights the major scattering effects of the gelatin medium on the OCT image quality (Supplementary Movie 1).

To quantify the quality of the OCT images acquired in different media (Fig. 1, F to H) we utilized the signal to noise ratio (SNR) formula 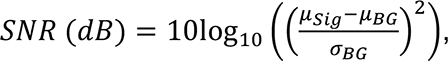 where μ_*Sig*_ is the average of the signal, μ_*BG*_ is the average of the background, and σ_*BG*_ is the standard deviation of the background (*31*). This analysis shows a linear decay in SNR with an average 6 dB higher SNR in water compared to embedded in gelatin and an average 9 dB higher SNR in water compared to embedded in the gelatin microparticle support bath (Fig. 1I). The SNR of the benchmark imaged with OCT in water (Fig. 1H) was sufficiently high to measure all channel widths using a custom MATLAB image analysis script and confirmed we printed the intended width for all channels (Fig. 1J). The 3D nature of the OCT images also allowed segmentation and 3D reconstruction of the printed benchmark model (Fig. 1K). The face-on and top-down views show the infill pattern as well as the channels. The isometric view of the bottom layers shows that the decay in SNR creates difficulties when segmenting the full print, resulting in an incomplete reconstruction. Together these data show the high performance of our custom 3D bioprinting platform and the potential of OCT for analyzing FRESH printed constructs, but that signal quality in the gelatin microparticle support bath must be significantly improved for in situ imaging.

### Image Quality is Improved with High Contrast Collagen and Sequential Imaging

As a next step to improve OCT image quality we incorporated titanium dioxide (TiO2) into the collagen bioink as a contrast agent (*32*). We selected TiO2 because it is a biologically safe compound for use in drugs and medical devices based on FDA guidance (*33, 34*). Printing the benchmark model using standard and high contrast collagen bioinks showed a clear improvement in image quality throughout the full depth (Fig. 2A), quantified as a 3-4 dB increase in SNR with the TiO2 (Fig. 2B). However, this still was not sufficient to image the entire benchmark model by OCT when embedded in the gelatin microparticle support bath.

**Figure 2.**
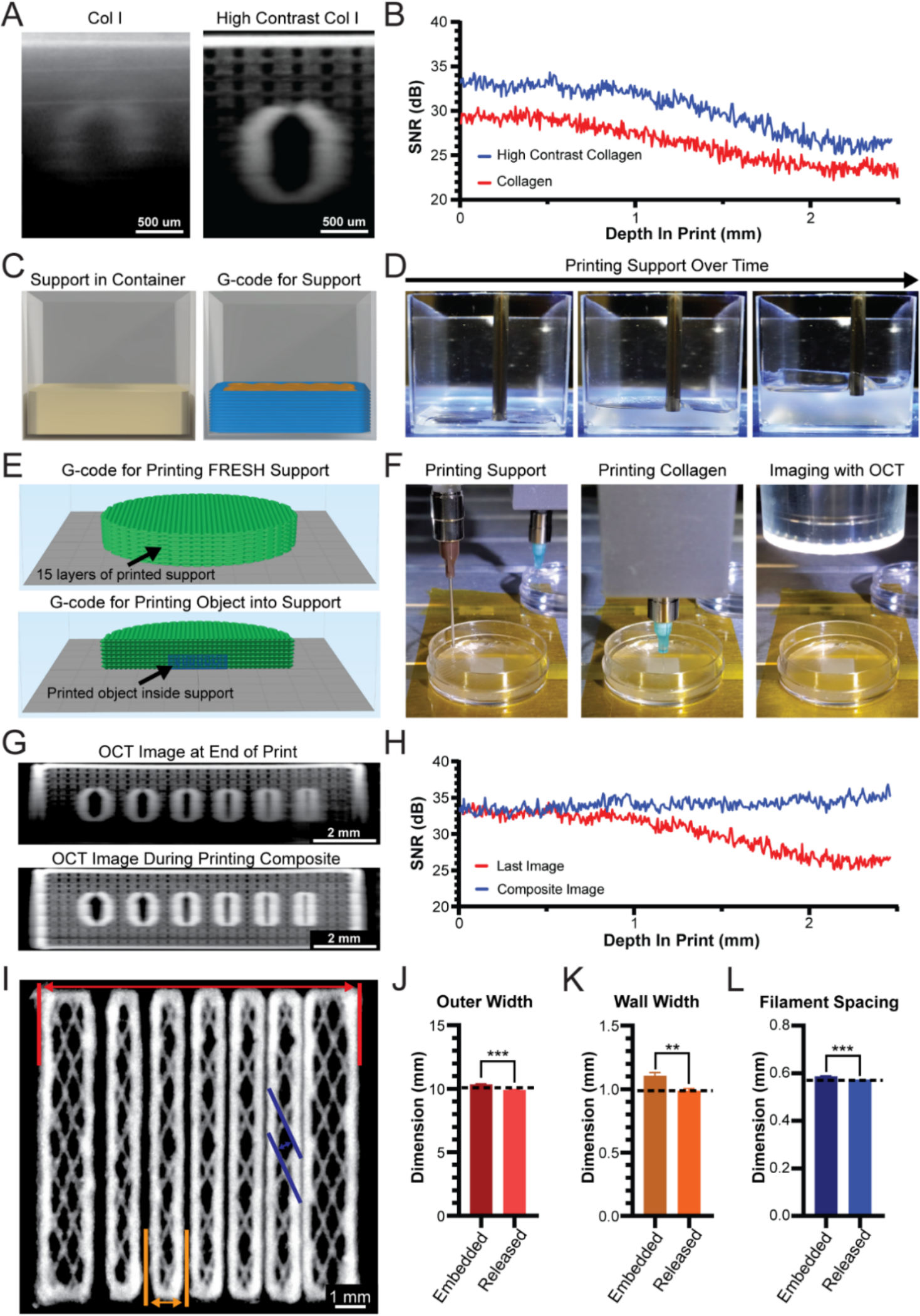
Development of a high contrast collagen bioink and printing of gelatin microparticle support bath to improve embedded OCT imaging. (**A**) OCT image cross-section of benchmark model channel printed from collagen-I or high contrast collagen-I embedded in the FRESH gelatin microparticle support bath. (**B**) OCT SNR analysis for a collagen and high contrast collagen benchmark model within the gelatin microparticle support bath. The two SNR curves are statistically significantly different from each other (p<0.0001, n = 9 total measurements from 3 benchmark prints each, Wilcoxon signed-rank test). (**C**) 3D render of FRESH gelatin microparticle support bath within a print container (left) and layer-by-layer G- code (right). (**D**) Time-lapse images of gelatin microparticle support bath printing. (**E**) G-code for printing the gelatin microparticle support bath (top) and for printing an object inside of the gelatin microparticle support bath (bottom). (**F**) Stepwise process of printing the support bath, printing a collagen construct inside of the support bath, and OCT image acquisition. (**G**) OCT imaging performed at the end of the print (top) or intermittently during printing to produce a composite image (bottom). (**H**) OCT SNR for image acquired at the end of print (Last Image) or by compositing in situ OCT images. The two SNR curves are statistically significantly different from each other (p<0.0001, n = 9 total measurements from 3 benchmark prints each, Wilcoxon signed-rank test). (**I**) Composite OCT cross-section image for evaluation of print fidelity in terms of outer width (red), channel wall width (gold), and filament spacing (blue). (**J**) OCT measured outer width while the print was embedded in the gelatin microparticle support bath or following print release (mean ± STD.; n = 3 prints, measurement at every x linescan in yz stack, embedded vs. CAD *P* = 0.0095, Released vs. CAD *P* = 0.0606, Embedded vs. Released *P* = 0.0004 [***] by Student’s two-tailed unpaired *t* test). Dashed line represents the 3D model expected value. (**K**) OCT measured wall width while the print was embedded in the gelatin microparticle support bath or following print release (mean ± STD.; n = 3 prints, highlighted wall measured at every x linescan in yz stack, embedded vs. CAD *P* = 0.0179, Released vs. CAD *P* = 0.3448, Embedded vs. Released *P* = 0.0026 [**] by Student’s two-tailed unpaired *t* test). (**L**) OCT measured filament spacing while the print was embedded in the gelatin microparticle support bath or following print release (mean ± STD.; n = 3 prints, 12 filaments each, embedded vs. CAD *P* = 0.0418, Released vs. CAD *P* = 0.0002, Embedded vs. Released *P* = 0.0008 [***] by Student’s two-tailed unpaired *t* test).

To further improve OCT image quality, we sought to decrease the scattering and absorption of the gelatin microparticle support bath by reducing the depth that must be imaged through. Instead of prefilling a container with the support bath and then FRESH printing within as we have previously done (*3–5*), we printed the gelatin microparticle support bath using a second printhead (Fig. 2C) that filled the container during printing (Fig. 2D). To do this we developed a custom MATLAB code that interleaves the G-code for the gelatin microparticle support bath and the G-code for the construct to be printed (Fig. 2E). In sequential steps, the gelatin microparticle support bath is first deposited, then a section of the construct is printed with high contrast collagen within the support bath, and then the OCT scan head is positioned over the print to acquire an image (Fig. 2F). Throughout the printing process OCT 3D image stacks of the printed construct are acquired at evenly spaced z-height intervals (Supplementary Movie 2). All acquired OCT stacks are then registered in 3D and stitched together to form a complete composite OCT 3D image stack of the entire printed construct. The composite OCT image shows a significant improvement in signal quality, specifically as depth increases (Fig. 2G, Supplementary Movie 3) (*35*). Importantly, we can see that for the composite OCT image the SNR no longer decays and by 2 mm in depth shows an improvement of >10 dB (Fig. 2H). Measuring external and internal features of our prints such as the overall size, width of the inner walls, and the spacing between filaments showed that the construct appeared as expected (Fig. 2I). Comparison of the construct in the gelatin microparticle support bath and after release confirmed that dimensions and print fidelity were maintained throughout the process (Fig. 2J-L and fig. S2). These results demonstrate the high precision and accuracy of our bioprinter and our ability to combine high contrast collagen I bioink, printing of the gelatin microparticle support bath, and sequential OCT capture for in situ 3D imaging and dimensional analysis.

### Image Quality is Improved by Increasing Transparency of the Gelatin Microparticle Support Bath

The light scattering by the FRESH gelatin microparticle support bath has always made it challenging to visualize constructs during the printing process. This stems from a difference in the refractive indices (RI) of the gelatin microparticles and the surrounding medium. To reduce scattering and improve transparency we needed to increase the RI of surrounding water (1.333) to match the RI of microparticles (Fig. 3A right) (*36, 37*). We initially replaced the aqueous buffer with a higher RI biologically compatible solution called histopaque (a mixture of polysucrose and sodium diatrizoate). However, the RI of the highest concentration histopaque (1.367) available was not sufficient to match the RI of the microparticles and only slightly improved transparency (data not shown). We next looked at histopaque components investigating high concentration polysucrose; however, at the concentrations necessary for RI >1.367, the solution viscosity increased considerably, and was incompatible with the yield stress behavior required for the FRESH gelatin microparticle support bath (*38*). We did not evaluate the other component of histopaque, sodium diatrizoate, because it is not iso-osmolar and therefore could result in cell damage.

**Figure 3.**
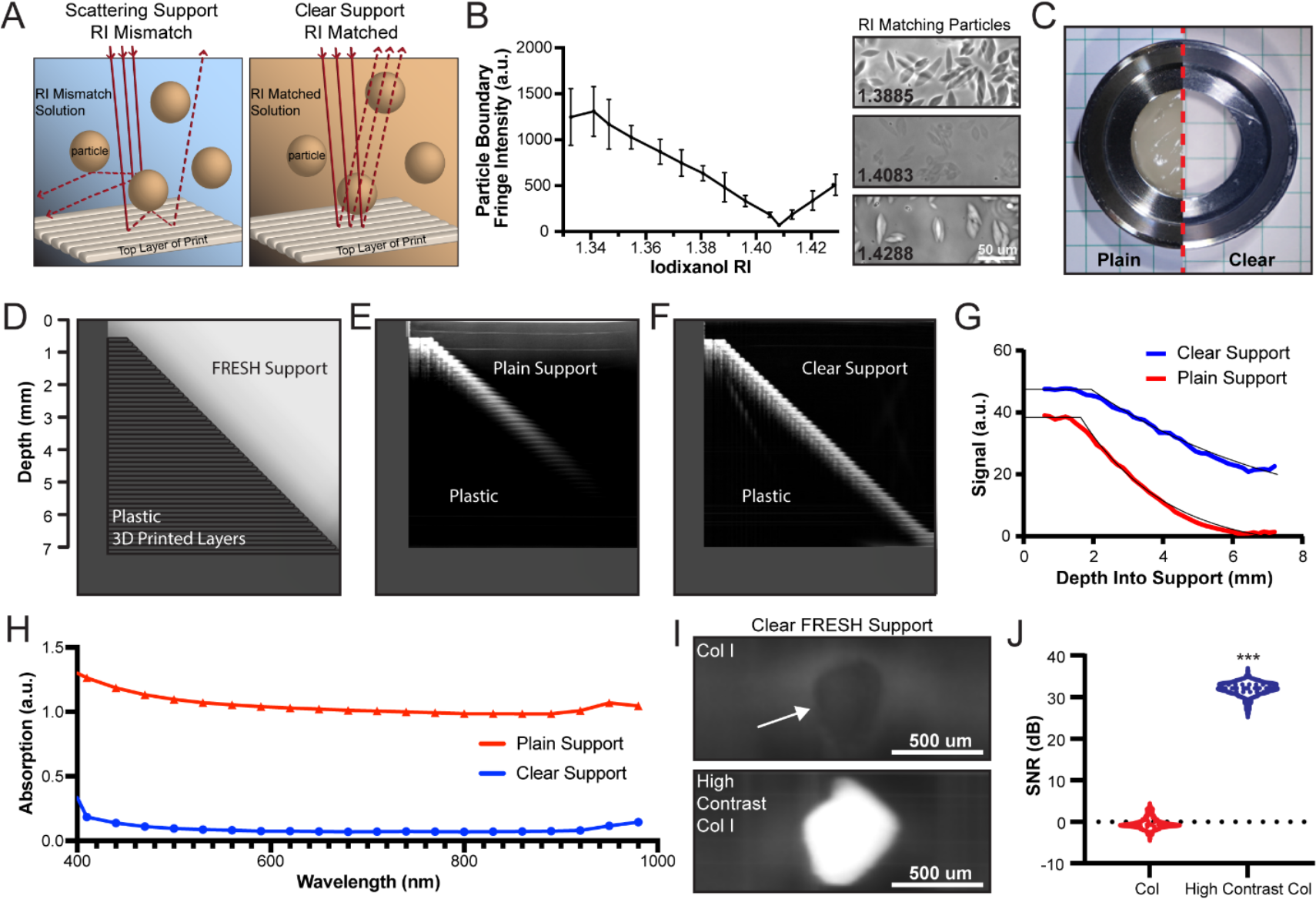
A transparent gelatin microparticle support bath increases OCT signal to noise and penetration depth. (**A**) For FRESH embedded printing, the gelatin microparticle support bath has an opaque appearance due to light scattering from the difference in RI between the particles and fluid phase solution. When the fluid phase RI matches the RI of the particles less light is scattered and absorbed which increases the gelatin microparticle support bath transparency. (**B**) Analysis of phase contrast imaging of gelatin microparticles’ boundary fringe intensity in solutions of iodixanol. Example images of gelatin microparticles in iodixanol at the RI inflection point. (**C**) Plain (left) vs the “clear” (right) gelatin microparticle support bath. (**D**) Depiction of a sloping 3D printed plastic well designed to determine the OCT penetration depth through gelatin microparticle support bath by visualizing the printed layers throughout the well depth. (**E**) OCT image through plain gelatin microparticle support bath. (**F**) OCT image through clear gelatin microparticle support bath. (**G**) Exponential fit and analysis of OCT signal penetration depth for plain (k = 0.4135) and clear gelatin microparticle support baths (k = 0.1709). There is a significant difference between both curves at all depths (p ≤ 0.0003, n = 4 containers each, two-way ANOVA with Tukey’s multiple pair- wise comparisons). (**H**) Absorption spectra for plain vs. clear gelatin microparticle support baths. Both curves are significantly different at all wavelengths (p < 0.0001, n=6 wells, 2-way ANOVA with Tukey’s multiple comparisons). (**I**) FRESH printed collagen-I (top) and high contrast collagen (Col)-I (bottom) filaments printed in clear gelatin microparticle support bath. (**J**) Analysis of the OCT SNR between collagen- I and high contrast collagen-I printed in clear gelatin microparticle support bath (mean ± STD.; n = 374 measurements along filament, *P* <0.0001 [***] measured by Student’s two-tailed unpaired *t* test).

Finally, we identified iodixanol as a non-ionic, iso-osmolar compound with high RI that is cell and tissue safe, endotoxin free, and comes as a 60% solution in water at a RI of 1.429 (*39*). To achieve a transparent FRESH support bath, we placed the gelatin microparticles within increasing concentrations of iodixanol with RI ranging from 1.333 to 1.429 and captured phase contrast images. By analyzing the fringe boundaries of the microparticles and the point at which they disappeared, we determined that they had an RI = 1.4083 (Fig. 3B and fig. S3). Preparing the microparticle gelatin support bath with iodixanol at this RI resulted in a highly transparent support, evident with a checkered background easily seen through a filled Petri dish (Fig. 3C).

To evaluate OCT performance in the clear gelatin microparticle support bath we 3D printed a plastic container with a sloping bottom to evaluate image quality at increasing depths (Fig. 3D). When imaging through the standard gelatin microparticle support bath the image quality degrades after a few millimeters, and the deepest parts of the dish cannot be resolved (Fig. 3E). In contrast, when imaging through the clear gelatin microparticle support bath the image quality is improved, and the deepest part of the dish can be resolved well (Fig. 3F, Supplementary Movie 4). Both support bathes display an exponential decay, with the signal in the plain support decaying faster (k = 0.4135) than in the clear gelatin microparticle support bath (k = 0.1709) (Fig. 3G). Absorbance spectral analysis of the clear support shows improved transparency throughout the visible-IR wavelengths (Fig. 3H) with an average decrease in absorption of 91.4 ± 2.3% (mean ± std) compared to plain support. Finally, within the clear support, the high contrast collagen bioink with TiO2 showed a significant increase in SNR compared to plain collagen bioink (Fig. 3I and 3J). This shows that by combining the clear support with the high contrast collagen bioink we can achieve high quality OCT imaging during the printing process (Supplementary Movie 5).

### In situ Volumetric Imaging and Fidelity Assessment of FRESH Printed Constructs

We next evaluated the combined effect of the clear gelatin microparticle support bath, high contrast collagen bioink, and sequential OCT imaging. In this case, using clear support means we did not need to print the support bath sequentially as it was sufficiently transparent for OCT imaging as is. Imaging the entire construct after printing showed some signal loss with imaging depth, however this was nearly eliminated by sequential OCT imaging to create a complete composite image (Fig. 4A, Supplementary Movie 6). Quantification confirmed this result, showing that the sequential OCT imaging minimized the linear decay in SNR observed when acquiring the OCT image only at print completion (Fig. 4B). This is qualitatively similar to the previous results with high contrast collagen bioink and printed layers of the gelatin microparticle support bath (Fig. 2H). However, using clear support we achieved substantially higher SNR values, resulting in further improvement in image quality. Finally, we confirmed that clearing with iodixanol did not affect the final printed dimensions or interfere with print release, with no noticeable changes in terms of print quality or fidelity (fig. S4).

**Figure 4.**
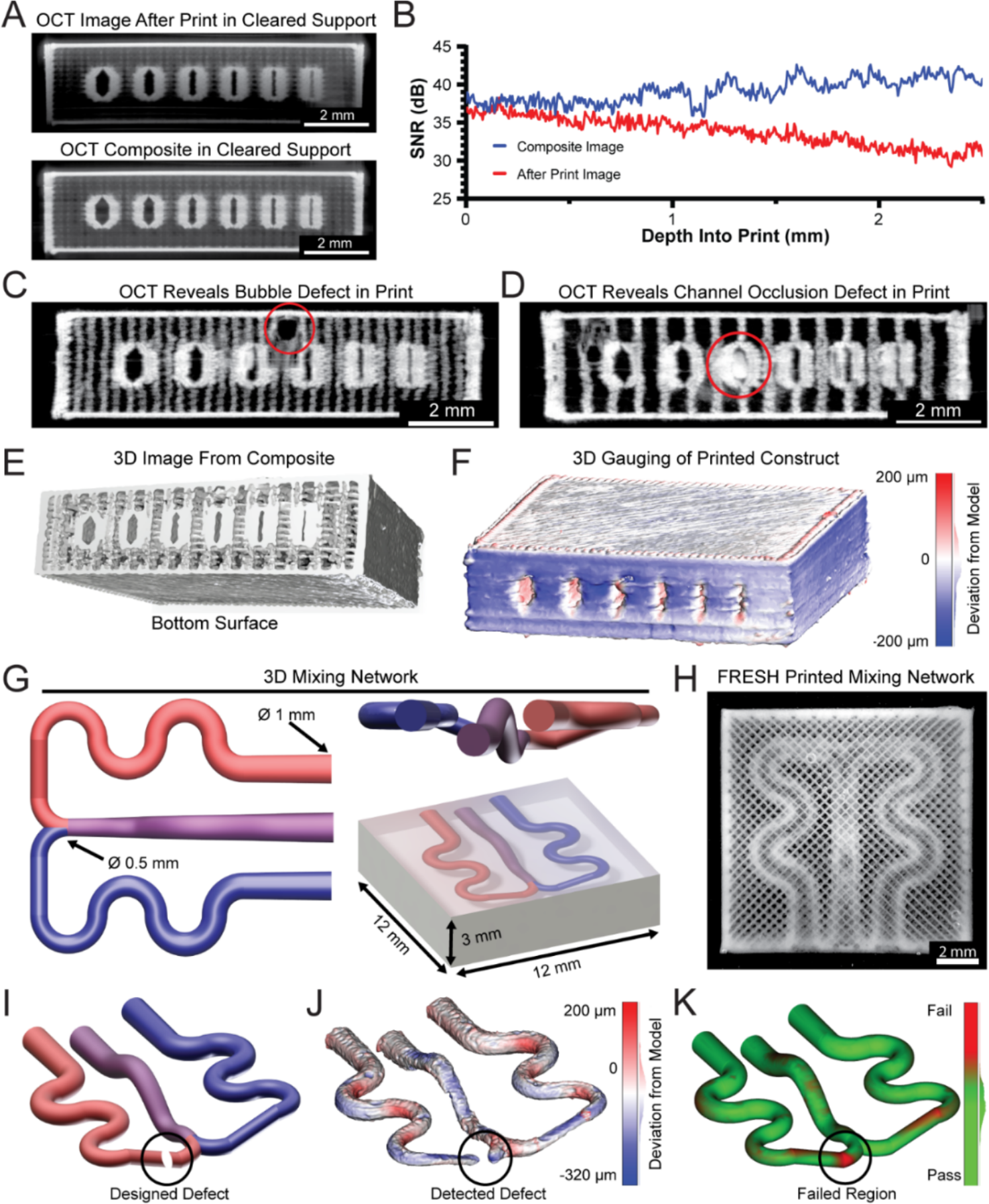
In situ OCT imaging and error detection of FRESH printed constructs. (**A**) Comparison of OCT imaging in clear gelatin microparticle support bath after print completion (top) or during in situ OCT imaging and composite image creation (bottom). (**B**) Analysis of OCT SNR after print imaging and during in situ imaging and composite image creation throughout the benchmark model depth. The two SNR curves are statistically significantly different from each other (p<0.0001, n = 9 total measurements from 3 benchmark prints each, Wilcoxon signed-rank test). (**C**) In situ OCT imaging reveals an internal print defect due to an introduced air bubble. (**D**) Unintentional channel occlusion detected by in situ OCT imaging. (**E**) 3D reconstruction using composite OCT image of a benchmark model. (**F**) Quantitative 3D gauging analysis to detect deviations in the reconstruction of the printed model compared to the computer-generated model. (G) 3D renders of a computer-generated mixing network to highlight FRESH printing and OCT capabilities. Top-down view of a FRESH printed 3D mixing network from collagen-I. (**I**) An engineered defect was designed into the mixing network to mimic a blockage to test the OCT detection capabilities. (**J**) Quantitative 3D gauging analysis revealing a void at the location of the engineered defect. (**K**) A pass-fail deviation analysis was implemented to compare the printed and imaged 3D volume to the expected computer model for validation or rejection of the final printed construct.

As an important next step towards quantitative assessment of print fidelity, the combination of high contrast collagen and clear gelatin microparticle support bath allows us to not only measure the quality of our prints, but also to detect errors. For example, we can identify internal void space errors, such as a bubble trapped in the top of the benchmark model during printing (Fig. 4C). In this case the sequential OCT imaging is critical, as most of the image below the bubble was captured before the bubble existed, which would cast a large shadow on features beneath it if the OCT image were acquired only at the end of the print. In addition to voids such as bubbles, we can also detect over extrusion errors, such as the blockage of a channel (Fig. 4D). Further, automatic image segmentation can be used to produce a full 3D reconstruction of the printed object followed by gauging software to measure deviations between the original 3D model and the 3D OCT reconstruction. We found an average deviation of –20.9 ± 60.4 µm, with most of this variation resulting from the external surface of the construct (Fig. 4E and 4F, fig. S5) (*40, 41*). While this 3D reconstruction was built using OCT images captured sequentially, time-lapse OCT imaging while the construct is being printed is also possible (fig. S6, Supplementary Movie 7). Such real-time imaging could allow for even more dynamic feedback on the printing process, allowing for error detection within a layer rather than waiting and imaging multiple layers.

In addition to basic defects, we can utilize in situ OCT imaging to systematically identify deviations in printed objects that would result in a loss of function. To demonstrate this, we designed a 3D microfluidic mixing network, with channels ranging in size from 0.5 to 1 mm within a collagen block (Fig. 4G). A photomicrograph of the mixing network 3D printed using high contrast collagen shows recapitulation of the intended geometry in a top-down view (Fig. 4H). To test our detection capabilities, we introduced a defect into the mixing network model that in practice could occur due to over-extrusion, filament dragging, or a pathing error, and would block flow from one side of the network rendering it non-functional (Fig. 4I, Supplementary Movie 8). After printing and imaging in the clear gelatin microparticle support bath we inverted the OCT images to extract the internal network shape and performed quantitative 3D gauging. Overall, the designed-defect mixing network showed high fidelity with an average deviation of –53.4 ± 62.1 µm, and clearly revealed the presence of the defect (Fig. 4J). By implementing a pass/fail criterion on the size of positive deviation we can automate the detection of this defect (Fig. 4K). While this analysis was performed after printing, it possible to implement this type of failure analysis during the printing process after each step of the sequential OCT imaging to provide a real-time error detection.

### In situ Imaging and Analysis of Complex 3D Human Tissue Constructs

Finally, we moved beyond the relatively simple benchmark model and mixing network to tissue constructs based on patient-specific medical imaging, leveraging our improvements in the bioink, support bath, and imaging. First, we printed a segment of the vestibular apparatus from the inner ear, which stress-tests the printing and imaging systems due to the patent semicircular canals oriented in three mutually orthogonal planes (Fig. 5A). The embedded print was clearly visible, and the optical imaging showed good reproduction of the model including patency of the three semicircular canals. The 3D OCT image also showed high signal quality, which allowed us to generate a 3D surface that revealed both external and internal features, with inner channels approximately 325 µm in diameter (Fig. 5A and Supplementary Movie 9). Gauging analysis showed that we printed with high-fidelity with an average deviation of 17.8 ± 43.3 µm. Second, we printed a segment of the circle of Willis from the brain’s arterial system, an unsupported patent vascular network that would be challenging to 3D bioprint without FRESH and would be almost impossible to image in an aqueous solution or air (Fig. 5B). Optical imaging of the embedded print showed good reproduction of the model including the vertebral arteries, the anterior spinal artery, the middle cerebral arteries, and the posterior and anterior cerebral arteries. The OCT composite image clearly showed the 3D structure, and automatic segmentation and 3D surface creation revealed the presence of both external and internal features, with the patent inner lumen of the right vertebral artery measuring approximately 450 µm in diameter (Fig. 5B and Supplementary Movie 10). Gauging analysis showed a vascular network with an average deviation of 21.8 ± 53.2 µm, comparable to the average deviation for the vestibular apparatus.

**Figure 5.**
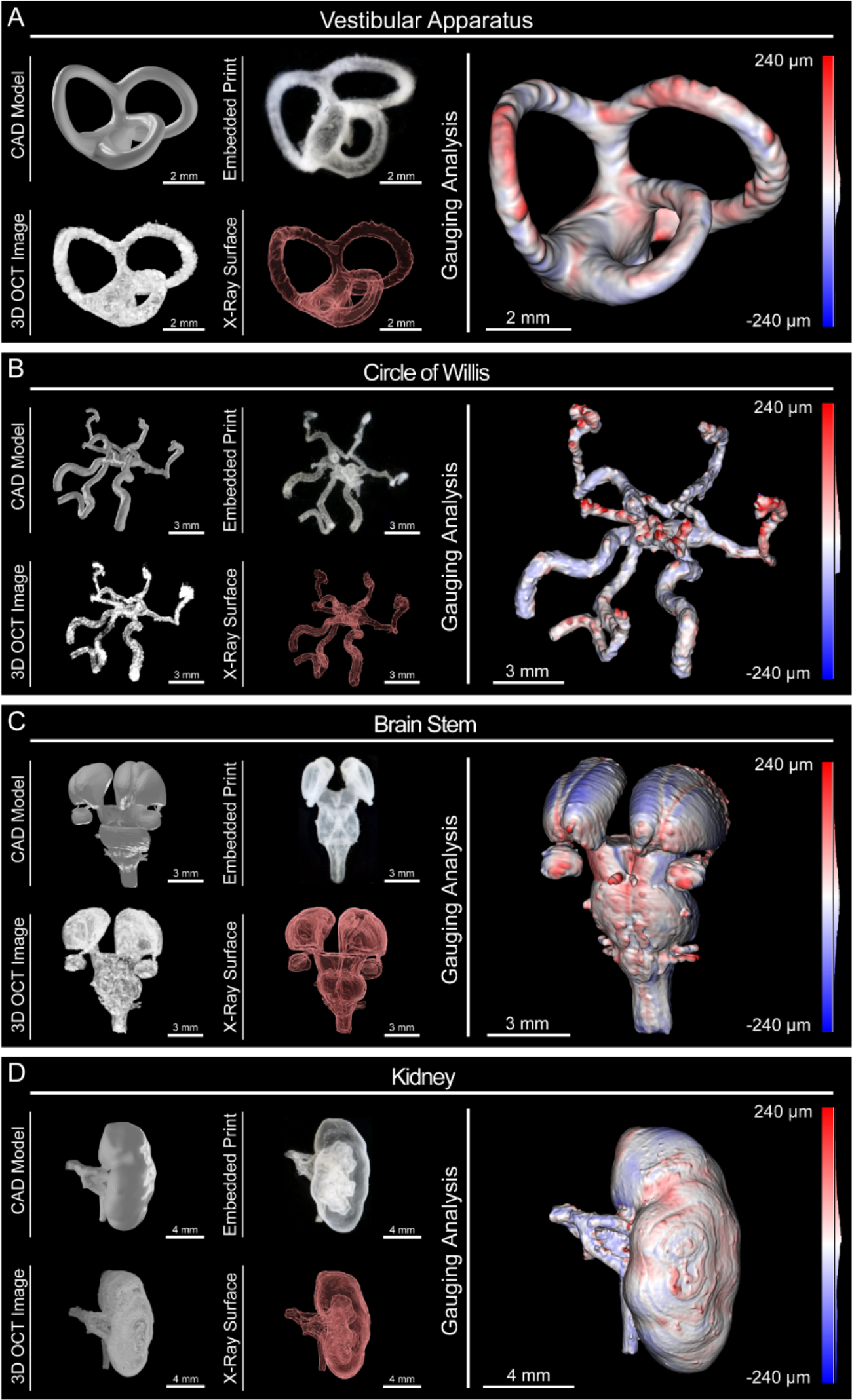
In situ OCT monitoring and 3D image reconstruction of medical imaging derived FRESH printed constructs. Four medical imaging derived models were chosen to demonstrate FRESH printing capabilities and OCT imaging using high contrast collagen in clear gelatin microparticle support bath. For each model we show five representations: 1) CAD model, 2) photograph of the embedded FRESH print in clear gelatin microparticle support bath, 3) an isometric view of the composite 3D OCT image, 4) an X-Ray view showing internal features, and 5) the results of quantitative gauging showing deviations of the 3D reconstruction from OCT imaging versus the computer model (red is oversized, blue is undersized). (**A**) Vestibular Apparatus with three orthogonal semicircular canals. (**B**) Circle of Willis showing vertebral arteries, anterior spinal artery, middle cerebral arteries, and posterior and anterior cerebral arteries. (**C**) Brain Stem with thalami, midbrain, pons, medulla and cranial nerves. (**D**) Kidney demonstrating internal calyces, ureter, and blood supply.

Next, we focused on tissue constructs based on solid organs, rather than tubular networks. The third model we printed was the brainstem, which is difficult due to the small size of the cranial nerves (∼250 µm in diameter) and their close proximity to each other (∼200 µm center to center) as well as the subtle surface features on the thalami (∼300 µm in depth) (Fig. 5C). Visually, the embedded construct was easy to see and displayed good reproduction of the CAD model including the thalami, the midbrain, the pons, and the medulla. The OCT composited image captured all the 3D structure and the 3D surface model revealed both external and internal features (Fig. 5C and Supplementary Movie 11). Gauging analysis revealed reproduction of the cranial nerves and subtle surface features of the thalami with an overall average deviation of 21.4 ± 63.1 µm, which is an order of magnitude below the smallest distinct feature within the model. The fourth model we printed was a scaled-down kidney to demonstrate the ability to print and image complex internal structures (renal artery, renal vein, calyces, and ureter) within a larger construct (the whole kidney) (Fig. 5D). The embedded print showed good reproduction of the CAD model with visible internal calyces as well as the renal artery, vein, and ureter. The OCT image confirmed high-fidelity printing throughout the depth of the kidney and the segmented 3D surface clearly showed external and internal features, including the open ureter, with an inner diameter of approximately 300 µm (Fig. 5D and Supplementary Movie 12). Gauging analysis confirmed recapitulation of the complex internal structures with an average deviation of -23.8 ± 121.9 µm. In total, the examples printed here highlight the ability to accurately print collagen tissue constructs from medical imaging data using FRESH bioprinting and validated using in situ OCT imaging.

## Discussion

One major challenge for 3D bioprinting has been monitoring the printing of soft and deformable hydrogels and creating a complete 3D reconstruction for dimensional validation and error detection (*6*). However, for 3D bioprinting to become a viable biomanufacturing platform, this limitation must be addressed to allow for the quality control and regulatory approval required for translation. For FRESH 3D bioprinting the gelatin microparticle support bath has limited visualization due to scattering and absorption by the particles themselves (*3, 4*). However, even when using transparent support baths (e.g. Carbopol), it has been difficult to use digital cameras to monitor print quality due to having to image through the support bath (*42, 43*). Many prints of interest have micron scale features, but overall dimensions on the order of tens of millimeters, which is a challenging range for many optical imaging techniques (*3–5*). Brightfield imaging can capture the entire printed object in one acquisition and be integrated into a 3D printer, but does not collect a 3D image. Other imaging modalities such as confocal and multiphoton microscopy have high resolution and optical sectioning for 3D reconstruction, but require long acquisition times and tile-scanning to capture larger volumes. Additionally, implementation of these techniques for in situ imaging would require either the construction of a custom confocal microscope or a shuttle system to transfer samples between the printer and microscope. Other 3D imaging techniques designed for acquisition of larger volumes such as µCT are capable of providing sufficient resolution, but typically require the incorporation of contrast agents as well as a long capture and reconstruction times that are not suitable for in situ imaging (*4*). For these reasons, we looked to OCT because it combines the large imaging volumes (cubic centimeters) and high-resolution (∼20 µm) of µCT, the optical sectioning of confocal imaging, and fast 3D acquisition times (5 seconds up to 3 minutes) similar to brightfield imaging.

Here we have shown that OCT can be effectively integrated into a 3D bioprinter system. The custom 3D bioprinter built here uses precision translation stages to provide easy mounting of the OCT scan head, the repeatable positioning (∼8 µm) for sequential imaging, and high printing speeds (up to 40 mm/s) (*44*). Commercial OCT scan heads are compact and light weight, with scanning mirrors, laser line and control inputs, and objective tightly packaged with height, width, and depth dimensions less than 10 cm. If the printing platform has sufficient space to mount this scan head, then incorporating OCT into a commercially available 3D bioprinter should be possible. With different printer configurations it would also be possible to have a stationary OCT scan head, to which the sample was moved (allowing for custom OCT systems), or beneath the sample for continuous live, in situ imaging (fig. S6, Supplementary Movie 7).

To date, FRESH and other embedded 3D bioprinting has used pre-filled containers of support material, whether the gelatin microparticle support bath or some other support bath such as alginate microparticles (*4, 6, 10*). However, for prints larger than ∼5 mm tall the deflection of the fine 34-gauge, 6.35 mm long tips limits resolution. We have previously overcome this by building custom needle tips with long, larger diameter and rigid tips terminated in small diameter tip (*5, 27*). However, printing into deep dishes of gelatin microparticle support bath presents other challenges such as dehydration and skinning of the upper layer of the support bath. Here we developed a new alternative approach by printing the gelatin microparticle support bath itself in order to minimize the deflection of the needle tip and without a limitation on construct height. In addition to high-resolution printing of larger constructs, this makes OCT imaging of an embedded print possible, as the thickness of support that must be imaged through is minimal (a few hundred microns vs. millimeters). A potential limitation of this technique could be the volume of the gelatin microparticle support bath syringe. Here we utilized a 10 mL syringe for our gelatin microparticle support bath, but a larger syringe could be adapted and used in a different configuration for greater support volumes.

OCT imaging of FRESH 3D biobioprinted constructs within the clear gelatin microparticle support bath provides a unique way to create 3D reconstructions without perturbing the soft and deformable hydrogel. Previously, to perform OCT imaging it was necessary to melt away the gelatin microparticle support bath and either submerge the construct in water or embed it in another clear hydrogel. In both cases, deviations in construct shape and size occurred due to the soft deformable nature of the hydrogel bioinks. Now, by using the integrated OCT and sequential imaging during printing we can create 3D reconstructions to directly gauge the printed object with the original digital model (*35, 41*). This type of gauging is important if 3D bioprinting is to become a mainstream manufacturing technology, and gauging is used extensively in subtractive manufacturing where coordinate measuring machines verify that the machined parts meet the required specifications (*45*). An important note about our approach is the use of automatic segmentation in the reconstruction of the 3D models, which prevents bias from manual intervention but can alter the results depending on the image SNR and algorithm used (*46*). In the future, machine learning algorithms could be trained on bioprinted constructs to improve automatic segmentation and provide higher accuracy when creating 3D reconstructions (*47*).

As 3D bioprinting advances there is a growing need for an in situ 3D imaging and validation system to determine dimensional accuracy and detect print errors. In all CAD and biologically-derived 3D models the FRESH printed constructs had average deviations of <50 µm be gauging analysis, demonstrating high-fidelity printing and sensitivity to micron scale error. This provides detection of surface deviations, but can also be used to perform more advanced pass/fail error detection including blocked channels, walls that are too thick or thin, defects caused by bubbles, and over extrusion of material. Indeed, in the future we intend to automate the routine analysis of 3D bioprinted constructs. For example, this would be useful in a microfluidic or vascular system where a hole could cause a leak, or for a tricuspid heart valve where fusion of the leaflets would block flow. These structural measurements of print quality are important because they impact functional performance and represent the type of quality control that will be necessary to comply with FDA and other regulatory agencies for clinical translation.

## Acknowledgments Funding

This work is supported by the Food & Drug Administration (R01FD006582), the Juvenile Diabetes Research Foundation (), and the National Heart, Lung, And Blood Institute of the National Institutes of Health (F32HL142229, 1F30HL154728, K99HL155777).

## Author contributions

All authors conceived the experiments and contributed to the scientific planning and discussions. J.W.T and D.J.S. prepared final figures and text. J.W.T conducted all bioprinting of constructs with in situ imaging. J.W.T and D.J.S. developed OCT imaging protocols and optical clearing protocols. J.W. T. wrote custom code in MATLAB for G-code editing. J. W.T and D. J. S performed image analysis in in FiJi and D.J.S. performed image analysis in Imaris. A.G. performed imaging and segmentation for creation of the kidney model. J.W.T, D.J.S., and A.W.F. wrote the paper and interpreted the data.

## Data and materials availability

All digital models, for the Replistruder 5 as well as printed and reconstructed models, are available for download at Zenodo.org **(ADD WEBSITE LINK)**. Any raw data not presented in the main and supplemental text is available on request.

## Materials and Methods

### Experimental Design

The objectives of the experiments presented here were to develop the ability to perform in situ imaging of FRESH bioprinted constructs using optical coherence tomography. To improve signal to noise ratio and image quality using OCT we experimented with high contrast collagen inks, bioprinting the gelatin microparticle support bath, and refractive index matching to make the support transparent. We then demonstrated the effectiveness of these approaches by imaging functional and dimensional deviations of multiple bioprinted constructs including four that were derived from medical imaging data.

### Integrated OCT Bioprinter

The bioprinter used here was built using four Parker Hannefin 404XR 100 mm travel precision stages in a gantry configuration (8 µm travel accuracy verified using Mitutoyo absolute digimatic indicator 543-792, data not shown) mounted to an aluminum baseplate (www.worldofclamping.com) (*44*). The printer utilizes two custom Replistruder 5 syringe pumps built with high-precision metric leadscrews (www.McMaster.com) and compact Nema 11 motors with planetary gearsets (www.stepperonline.com); both were mounted to the Z stage. A dovetail adapter was designed, 3D printed from PLA plastic, and mounted to the Z stage to receive the OCT scanhead (Thorlabs), which was controlled with its own dedicated PC. The X, Y, and Z axes as well as the two extruders were controlled using a Duet 3 motion controller with a Raspberry Pi 4 single board computer dedicated for the user interface.

### Plain FRESH Gelatin Microparticle Support Bath and Generation

FRESH v2.0 gelatin microparticle support bathwas prepared as previously described using a complex coacervation method to produce gelatin microparticles (*4*). First, 2.0% (w/v) gelatin Type B (Fisher Chemical), 0.25% (w/v) Pluronic® F-127 (Sigma-Aldrich) and 0.1% (w/v) gum arabic (Sigma-Aldrich) were dissolved in a 50% (v/v) ethanol solution at 45°C in a 1 L beaker and adjusted to 7.5 pH by addition of 1M hydrochloric acid (HCl). The beaker was then placed under an overhead stirrer (IKA, Model RW20), sealed with parafilm to minimize evaporation, and allowed to cool to room temperature while stirring overnight. The resulting gelatin microparticle support bath was transferred into 250 mL containers and centrifuged at 300 g for 2 min to compact the gelatin microparticles. The supernatant was removed and gelatin microparticles were resuspended in a 50% ethanol solution of 50 mM 4-(2-hydroxyethyl)-1-piperazineethanesulfonic acid (HEPES) (Corning) at pH 7.4, to remove the Pluronic® F-127. The gelatin microparticle support bath was then washed three times with the same Ethanol HEPES solution and stored until use at 4C. Prior to printing, the uncompacted support was centrifuged at 300 g for 2 minutes then washed with 50 mM HEPES and centrifuged at 750 g for 3 minutes a total of 4 additional times. After the last washing the gelatin microparticle support bath was again suspended in 50 mM HEPES and was degassed in a vacuum chamber for 15 min, followed by centrifugation at 1900-2100 g, depending on level of compaction desired, for 5 min. The supernatant was removed and the gelatin microparticle support bath was transferred into a print container.

### Collagen Bioink Preparation

All collagen bioinks were purchased as LifeInk 200 (Advanced Biomatrix). For bioprinting these inks were prepared as previously described (*4*). Briefly, 35 mg/mL LifeInk was mixed with syringes in a 2:1 ratio with .24M acetic acid to produce a 23.33 mg/mL acidified collagen ink. The ink was then centrifuged at 3000 g for 5 minutes to remove bubbles. To produce high contrast collagen inks an appropriate amount of 0.3-1.0 µm TiO2 powder (Atlantic Equipment Engineers) was then weighed out for a 250 PPM mixture with the acidified collagen bioink. The TiO2 powder was then dissolved in 100 µL of .24M acetic acid. This TiO2 solution was then aspirated into the collagen through a needle. The TiO2 collagen mixture was then mixed 100 times between two syringes. The ink was then centrifuged at 3000 g for 5 minutes to remove bubbles. For printing the bioinks were transferred to a 500 µL gastight syringe (Hamilton Company).

### OCT Imaging

To acquire an image with the Thorlabs Vega 1300 nm OCT system (VEG210C1) the sample is first placed under the objective (OCT-LK4 objective). The system was started in the 2D mode with a scanline intersecting the sample. The sample’s surface was then brought into focus and shifted using the reference stage to highlight the region of interest and to set further parameters. The amplification and reference intensity were then set to provide the highest signal without introducing image artifacts. The polarizing filters were then adjusted to optimize the signal intensity and minimize image artifacts. For a 2D image the averaging and z depth were then set, and the image was acquired at this point. For a 3D volume the mode was switched to 3D then the x, and y pixel dimensions were set to provide sufficient resolution while allowing for averaging and the amount of averaging was set (typically 16.22 µm or 20 µm with 10 averages). Finally, the image was acquired. When utilizing the OCT mounted to the bioprinter for in situ imaging this same process was executed after the printer automatically positioned the scan head and paused for imaging. Acquired images were exported as 32-bit Tiff files for further processing.

### Printing Both Bioink and Gelatin Microparticle Support Bath with In situ Imaging

To generate print pathing for multimaterial printing with in situ imaging we used a combination of open-source software and custom code. First the object to be printed and the volume of gelatin microparticle support bath to be printed were generated using Autodesk Inventor (Autodesk) or acquired from another source. These models were then loaded into Ultimaker Cura (Ultimaker) and processed into G-code using print parameters appropriate to the needle and syringe diameter being used. Next the G-codes were imported into a custom MATLAB script (Mathworks) designed to interleave the support print, the collagen print, and the imaging steps (meshGcode_OCTandSupport.m). The script takes advantage of the ability built into the Duet 3 implementation of G-code to store multiple toolhead positions. Using an index of the layer change comments in the G-code (automatically generated by Cura) the MATLAB script inserts small G-code scripts that use these toolhead positions to swap between extruder 1 (bioink), extruder 2 (gelatin microparticle support bath), and the OCT scan head. This custom script also allows for selection of the number of initial layers of gelatin microparticle support bath prior to initializing the collagen print and the number of layers of collagen to print prior to a new layer of support. The OCT scan head is also automatically raised by the thickness of the new layers printed to maintain focus on them. Prior to printing, the high contrast collagen bioink is centrifuged at 3000g for 5 minutes in a 10 mL plastic BD syringe and 450 µL is transferred to a 500 µL Hamilton gastight syringe. The plain gelatin microparticle support bath is centrifuged at 2000 g for 5 minutes in a 10 mL plastic BD syringe and is transferred to a 10 mL Hamilton gastight syringe. These inks, in their syringes, are loaded into their dedicated Replistruder 5 syringe pumps. Then, using the tip of the needle to measure the width and height, the first needle is aligned to the center of the print dish. For this first tool the origin is set using the G92 command. The first toolhead (T0) position is then set using the G10 command. Next, the second extruder’s needle is centered on the dish and the position is again recorded using the G10 command, but for the second toolhead (T1). Finally, the OCT is aligned by centering its objective on the dish. Next the Z positions must be set. The first extruder (T0) is touched off to the bottom surface of the print dish and the G92 and G10 commands are used to set its Z offset. This process is repeated with the second extruder (T1) and the G10 command. Finally, the OCT is shifted in Z until the boundary at the bottom of the dish is in focus and the G10 command is used to set the offset of the focal plane. At this point the Duet 3 knows all the relative positions of the tools attached to the Z axis and is ready to print.

The second extruder (T1) is then returned to the center of the dish, using the measured offsets from the alignment process. Plastic dishes are filled with DI (for collagen bioink) or 50 mM HEPES (for support ink) and placed such that the inactive extruder needle is submerged (to prevent drying out and clogging). At this point the print can be initiated by selecting the G-code on the Duet 3 interface. When the printer reaches the first pause for imaging the user can acquire an image using the Vega OCT’s PC and then reinitiate printing using the Duet 3 interface.

### Index of Refraction Measurement

Index of refraction measurements for Iodixanol solutions were taken using a Hanna Instruments digital refractometer (HI96800). Briefly the refractometer was calibrated using deionized water, then the sample of interest was placed on the flint glass for measurement. The sample was allowed to equilibrate in temperature with the steel ring of the refractometer. Measurements were repeated until the 4^th^ digit was consistent across measurements.

### Identification of Gelatin Microparticle Support Bath Index of Refraction

Plain gelatin microparticle support bath was prepared as described above. A high viscosity pipette was used to transfer 100 uL of compacted gelatin microparticle support bath into 2 mL of prepared Iodixanol solutions ranging from 0% to 60% in a 12 well plate. The support was then dispersed thoroughly within the Iodixanol solutions using clean pipette tips. Coverslips were used to trap the particles against the bottom of the well, which was necessary as Iodixanol solutions above 30% were denser than the gelatin microparticle support bath and floated the particles. Images were then acquired using a Nikon Eclipse TS100 microscope using a 20X phase objective and a Photometrics CoolSnapES camera run by MicroManager. The images were captured with identical exposure and illumination settings to allow for direct comparison.

The Iodixanol solutions were measured with our digital refractometer as described previously. To compare the particle clarity at different indices of refraction the peak to trough difference of the high contrast phase boundary was quantified using a line scan analysis in FIJI ImageJ software for n = 13 particles in each image. Then the absolute value of this was plotted against the refractive index to identify the refractive index and iodixanol concentration that resulted in the clearest support solution.

### Clear FRESH Gelatin Microparticle Support Bath Generation

To produce clear gelatin microparticle support bath using iodixanol the initial process is the same as plain support, described above except 250 mM HEPES is used instead of 50 mM HEPES. After the plain support is prepared for printing (e.g., centrifuged at 2000g for 5 minutes and supernatant removed), the supernatant is poured out and the remaining supernatant is absorbed with Kim wipes. The desired final concentration of Iodixanol ranged from 47.5% to 50% for optimal clarity and print characteristics, here we demonstrate bringing the concentration to 50%. First, Iodixanol reused from previous support preparations, at roughly 50%, is measured using our Hanna Instruments digital refractometer. The refractive index is utilized to determine the exact Iodixanol concentration. Sufficient Iodixanol is added to the support to bring the mass ratio of Iodixanol to 26% (mass ratio of 1:1.1 support:∼50% Iodixanol). The solution is then vortexed. After mixing the 26% Iodixanol support is centrifuged at 3500 g for 5 minutes. After compaction, the Iodixanol supernatant is removed. At this point new 60% Iodixanol is added at a mass ratio sufficient to bring the final solution up to 50% Iodixanol (mass ratio of 1:2.3 support:60% Iodixanol). This 50% Iodixanol support solution is backfilled into capped 10 mL BD syringes (with plungers removed) and placed in a vacuum chamber for 15 minutes. The open barrel of the syringe was sealed with parafilm and then the syringes were centrifuged at 3500 g for 15 minutes. After centrifugation, the denser Iodixanol is on the luer lock side of the syringe, covered by the compacted, clear gelatin microparticle support bath. A plunger is reintroduced into the backside of the syringe using a thin wire to allow for the passage of air. The Iodixanol solution is removed to another 10 mL BD syringe using a luer coupler and collected for later use in the first Iodixanol wash. After the Iodixanol has been removed, the clear gelatin microparticle support bath is collected into as few BD syringes as can hold the total volume. The support is transferred back and forth 50 times between these syringes to homogenize the mixture. Finally, the support is transferred to new 10 mL BD syringes with their plungers removed. The open barrel is covered with parafilm and the syringes are centrifuged at 3500g. The plungers are reinserted, again using a wire to break the seal, and the support is ready for use.

### Absorbance Spectra Measurement

To measure the absorbance of different gelatin microparticle support bath preparations we used a spectrophotometer (Molecular Devices SpectraMax i3x). For each preparation, an equal volume of gelatin microparticle support bath was deposited into the wells of a standard clear 24 well plate (thus producing an equal thickness). The spectrophotometer was set to acquire absorbance spectra from 230 nm to 980 nm in 30 nm steps for each filled well. The plastic dish absorbed light from 230 – 400 nm and so this range was excluded from the analysis. Measurements were taken at a consistent temperature of 24°C.

### Printing in Clear Gelatin Microparticle Support Bath with In situ Imaging

Generating print pathing for printing in clear gelatin microparticle support bath with in situ imaging is very similar to printing both the bioink and the support. Ultimaker Cura is utilized to generate the print pathing for the object to be printed. Next the G-code is imported into a custom MATLAB script (Mathworks) designed to interleave the collagen print and the imaging steps (meshGcode_OCT.m). The resulting G-code is uploaded to the Duet 3 printerboard via USB stick.

Prior to printing, the dish is filled with clear gelatin microparticle support bath and the top surface is scraped flat using a 20 mm x 20 mm square 1.5 coverslip. The top surface of the support is then covered in a layer of light mineral oil to prevent drying out (Fisher Scientific). Next the offset between the bioink toolhead (T0) and the OCT scan head (T2) is measured as previously described and stored using the G10 tool offset command. The bioink extruder (T0) is then returned to the center of the dish, using the measured offsets from the alignment process. A plastic dish is filled with DI (for collagen bioink) and placed such that the inactive extruder needle is submerged (to prevent drying out and clogging during imaging). At this point the print can be initiated by selecting the correct G-code on the Duet 3 interface. When the printer reaches the first pause for imaging the user can acquire an image using the Vega OCT’s PC and then reinitiate printing using the Duet 3 interface.

### 3D Model Creation

All models were created using Inventor Professional 2020 (Autodesk) or downloaded from online repositories. For the benchmark model and the 3D mixing model, the entire model was generated in Inventor then exported as an STL for printing. The vestibular apparatus was sourced from https://vestibularfirst.com/how-to-print-3d-vestibular-apparatus/ and was derived from MRI data acquired by the University of Dundee School of Medicine (https://sketchfab.com/3d-models/anatomy-of-the-inner-ear-f80bda64666c4b8aaac8f63b7b82a0a0). The circle of Willis, derived from MRI data, was sourced from the NIH 3D Print Exchange (Model ID 3DPX-002604). The brain stem model, derived from MRI data, was sourced from the NIH 3D Print Exchange (Model ID 3DPX-003892). The kidney model was sourced from the University of Pittsburgh School of Medicine.

### Image Analysis

To assess signal to noise ratio in OCT images they were first opened in FIJI ImageJ. Then areas of the image with high signal and adjacent areas of background were sampled. The resulting averages and standard deviations for each area were utilized in the equation 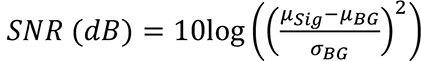 to calculate the signal to noise ratio, where μ_’()_ is the average of the signal, μ_*+_ is the average of the background, and σ_*+_ is the standard deviation of the background (*31*).

To assess the signal intensity at different depths in plain and clear gelatin microparticle support bath a plastic dish was created with a sloped bottom that allowed for imaging through increasing depths of gelatin microparticle support bath. Images of the bottom of the dish were acquired using plain and clear gelatin microparticle support bath at the same imaging settings. Images were opened in FIJI and converted to 16-bit TIFFs using the same range of intensity values prior to conversion. The images were scaled to match each other, then regions of the same size were sampled at equivalent depths in each image. The resulting averages were utilized to create a curve of intensity at depth for plain and clear gelatin microparticle support bath.

To assess the quality of the printed resolution test object the acquired OCT images were opened in FIJI. The images were corrected for rotation about the x, y, and z axes. Cropped views of the middle 50% along the channel axis and 25% perpendicular to it were extracted from the images. These images were then 3D-median filtered with a 5- pixel filter width. The background intensity in the center of the channel was measured and used to perform background subtraction. The images were then automatically thresholded. The images were cropped to only include the channels and their walls. These stacks were saved and then imported into a custom MATLAB script that measured the width of the channel at each row in each slice of the segmented images and reported the average value of this width for all slices.

### OCT Image Composites

To combine multiple OCT image stacks acquired throughout printing into a single composite image the pairwise stitching plugin was used in FIJI ImageJ with subpixel accuracy and linear gradient or maximum intensity fusion options set (*35*). First the raw images were rotated to orient the print to the pixel x, y, and z axes. Next the images were cropped to include the full last layer and the bottom of the print. These images were then resliced to a view perpendicular to the Z axis. The first stitching was performed with the first and second captured OCT stacks. The second stack was cropped to have approximately 10% overlap with the first. Overlapping regions of interest were highlighted to aid the plugin in registration. In subsequent stitching steps the same process was repeated but using the previously stitched image and the next stack to be incorporated, until the last stack was incorporated.

### Gauging

To perform quantitative gauging of printed objects using the OCT images a full model of the printed object needed to be extracted. To do this the OCT images, either one acquired at the end of printing or a composite of multiple acquired throughout printing, were loaded into FIJI (NIH). These stacks were then filtered using the 3D-Median filter with a filter width of 3 pixels. The background subtraction tool was then used to remove background if necessary. The images were then converted into 16-bit tiffs and the histogram equalization was used. Local automatic thresholding was utilized to isolate the image data, using the savola method. Next erroneous spots were removed using the outlier removal tool. Finally, the segmented image stack was exported as a tiff.

The tiff was imported into 3D Slicer (http://www.slicer.org/). The integrated segmentation tools were used to isolate the print. The segmentation was manually edited to remove artifacts and islands and was then smoothed using a median filter with a 3 pixel width. After smoothing the reconstruction was exported as an STL file using the built-in exporter, taking care to set the pixel size to match the known pixel dimensions from OCT acquisition.

The reconstruction was loaded into 3D builder (Microsoft), where scaling was verified, and the mesh was simplified to aid with future processing. After saving the simplified mesh the reconstructed object was loaded, along with the digital model STL (which was used to generate the G-code for printing the object) into CloudCompare (http://www.cloudcompare.org/) (*41*). Using built in tools the two objects were oriented relative to each other, registered, then surface deviations were calculated. This information was exported as false color images as well as mean and standard deviation.

### 3D Visualization in Imaris

We used Imaris (Bitplane, 9.5.1) for 3D rendering of the raw OCT data. The TIF file from the OCT data for each organic model was imported into Imaris. A surface object was created using the surface wizard function with local background subtraction and filtered using the “quality” filter to remove small non-specific objects. A mask of the surface object was created to act as a passthrough filter for the original OCT data to remove non-specific background from the 3D image. 3D renders of the background removed OCT data and the X-ray view of the surface object were exported as TIF images. Built in animation functionality was then used to make movies showing the internal features and highlighting internal complexity within the printed objects.

### Statistics and Data Analysis

Statistical and graphical analyses were performed using Prism 9 (GraphPad) software and Excel (Microsoft v16). Statistical tests were chosen based on the experimental sample size, distribution, and data requirements. For comparison of SNR between benchmark models in plain gelatin microparticle support bath, gelatin, and water Wilcoxon paired signed-rank tests were used (Fig. 1I). For analysis of measured width of a benchmark imaged in water the Pearson correlation coefficient was calculated (Fig. 1J). For comparison of SNR between benchmark models in plain support using plain and high contrast collagen Wilcoxon paired signed-rank tests were used (Fig. 2B). For comparison of SNR between last image and composite image of benchmark models in plain support using high contrast collagen Wilcoxon paired signed-rank tests were used (Fig. 2H). For comparison of embedded and released dimensions of benchmarks printed in plain support Student’s two-tailed unpaired t test was used (Fig. 2 J to L). For comparison of OCT signal in plain and clear gelatin microparticle support bath at all depths 2-way ANOVA with Tukey’s multiple comparisons was used (Fig. 3G). For comparison of absorbance in plain and clear gelatin microparticle support bath at all depths 2-way ANOVA with Tukey’s multiple comparisons was used (Fig. 3H). For comparison of SNR of plain collagen and high contrast collagen in clear gelatin microparticle support bath Student’s two-tailed unpaired t test was used (Fig. 3J). For comparison of SNR between last image and composite image of benchmark models in clear gelatin microparticle support bath using high contrast collagen Wilcoxon paired signed-rank tests were used (Fig. 4B). For analysis of measured width of a benchmark printed in printed support after release the Pearson correlation coefficient was calculated (fig. S2B). For analysis of measured width of a benchmark printed in clear gelatin microparticle support bath after release the Pearson correlation coefficient was calculated (fig. S4B). Preparation of figures and visuals was completed in Adobe Photoshop and Illustrator CS6 and CC. OCT images were edited in Fiji (ImageJ NIH) and Imaris 9.5.1 (Bitplane). 3D Reconstruction of OCT images was completed using 3D Slicer (http://www.slicer.org/). Gauging was accomplished using CloudCompare (http://www.cloudcompare.org/). Advanced image analysis and quantification was performed in MATLAB (Mathworks).

