## Supplemental Figures for "In Situ Volumetric Imaging and Analysis of FRESH 3D Bioprinted Constructs Using Optical Coherence Tomography"

**Title**

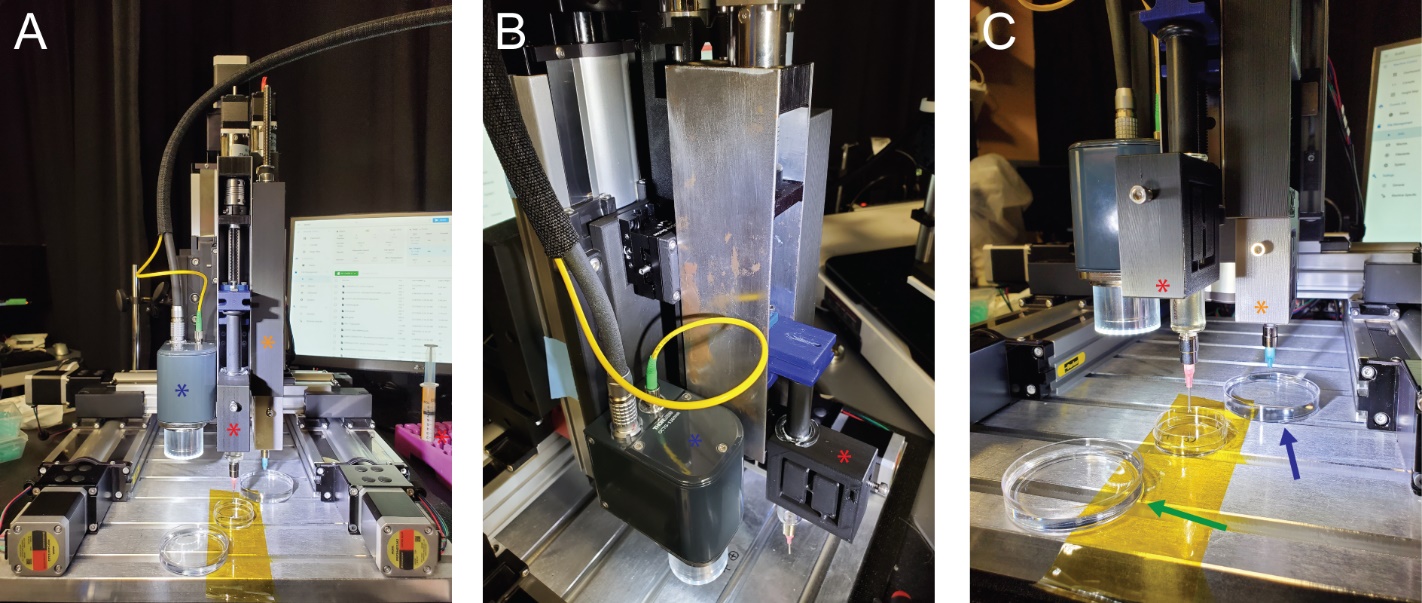

**Figure S1| Photographs of custom 3D bioprinter with OCT scan head mounted.** (**A**) Front on view showing the gantry configuration of the printer with OCT scan head on the left (blue *) and two Replistruder 5 syringe pumps on the right (red and orange *). (**B**) Angled view showing the z stage, a closer view of the OCT scan head (blue*) and the Replistruder 5 assigned to gelatin microparticle support printing (red *). (**C**) Right side view of the printer set up for printing high contrast collagen I in printed plain gelatin support. The left syringe (red *) is filled with gelatin microparticle support bath and the right with high contrast collagen I (orange *). The dishes diagonally aligned are filled with 50 mM HEPES (left, green arrow) and DI water (right, blue arrow).

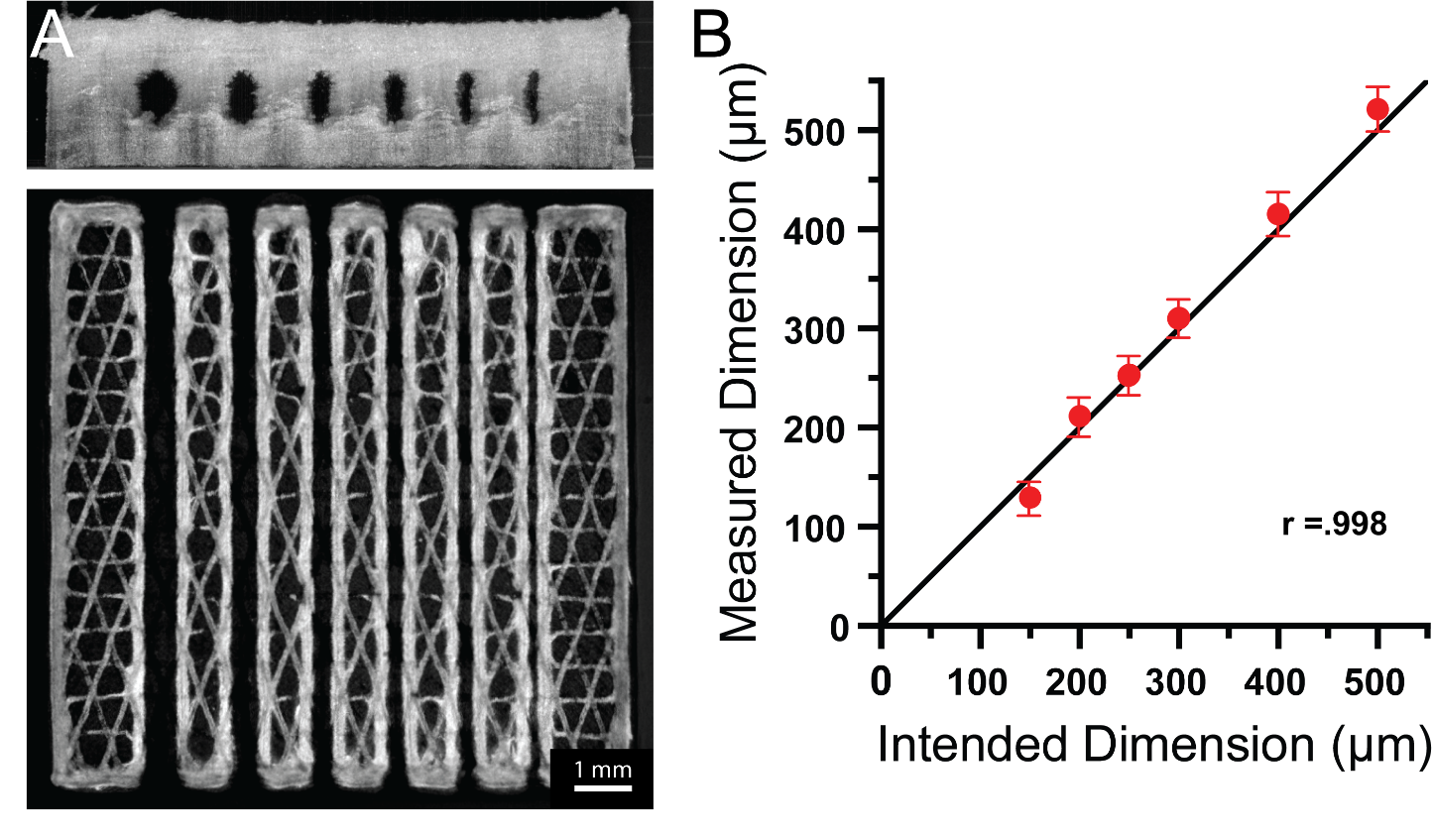

**Figure S2| High contrast collagen print released from printed plain gelatin support retains its shape.** (**A**) An end on (above) and top down (below) view of a benchmark model printed using high contrast collagen I in printed plain gelatin microparticle support, after melting away the support. (**B**) Measured versus intended dimensions for the released print RMS error = 14.9951 µm (mean ± STD; n = 1 print with 12978 measurements, Pearson correlation coefficient = .998).

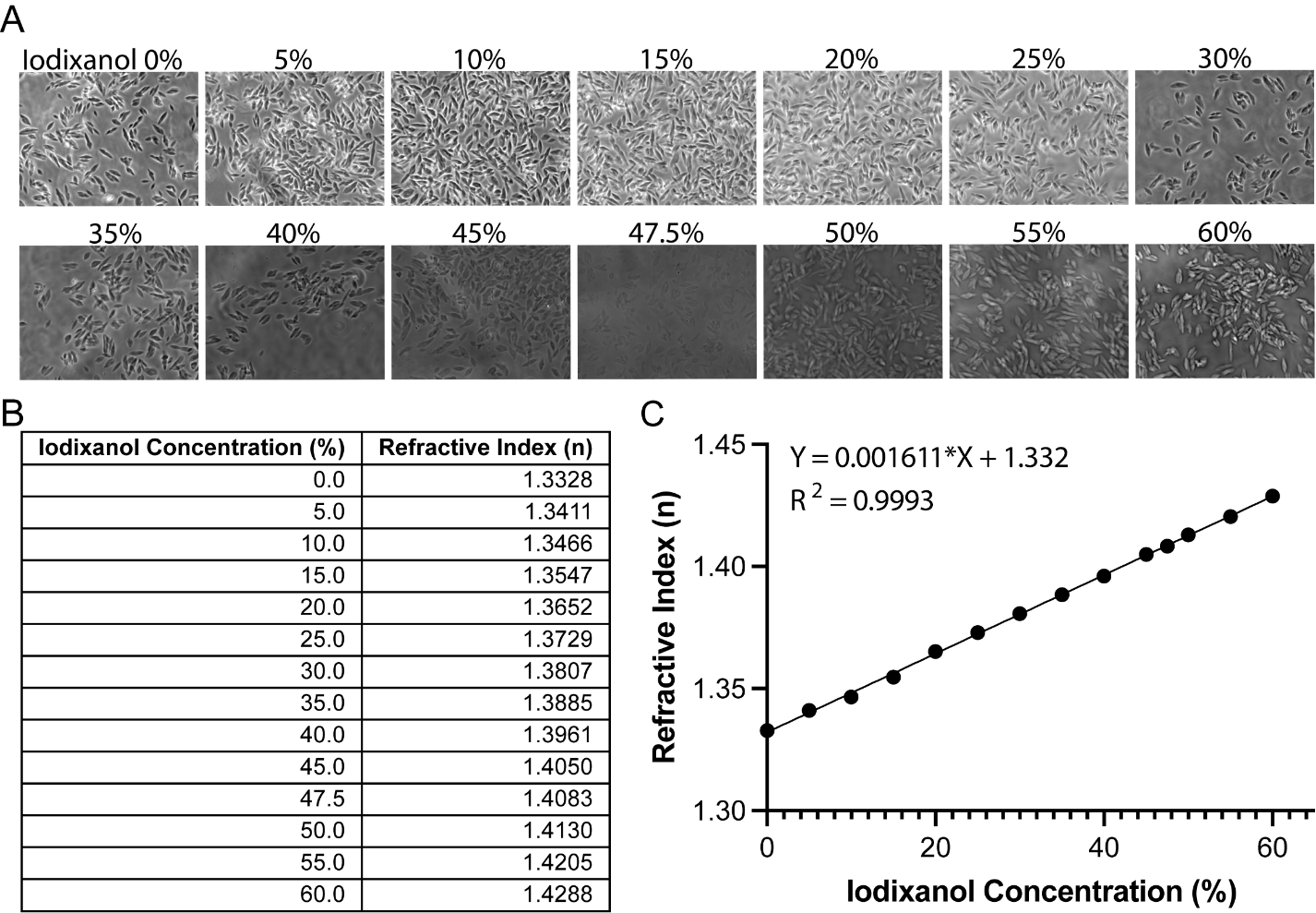

**Figure S3| Sweep of iodixanol concentration to identify gelatin microparticle support refractive Index.** (**A**) Gelatin microparticles were suspended in concentrations of iodixanol varying from 0% to 60%, with corresponding refractive indices of 1.333 and 1.429. (**B**) Table of Iodixanol concentrations with corresponding measured refractive index. (**C**) Refractive index versus iodixanol concentration with linear fit.

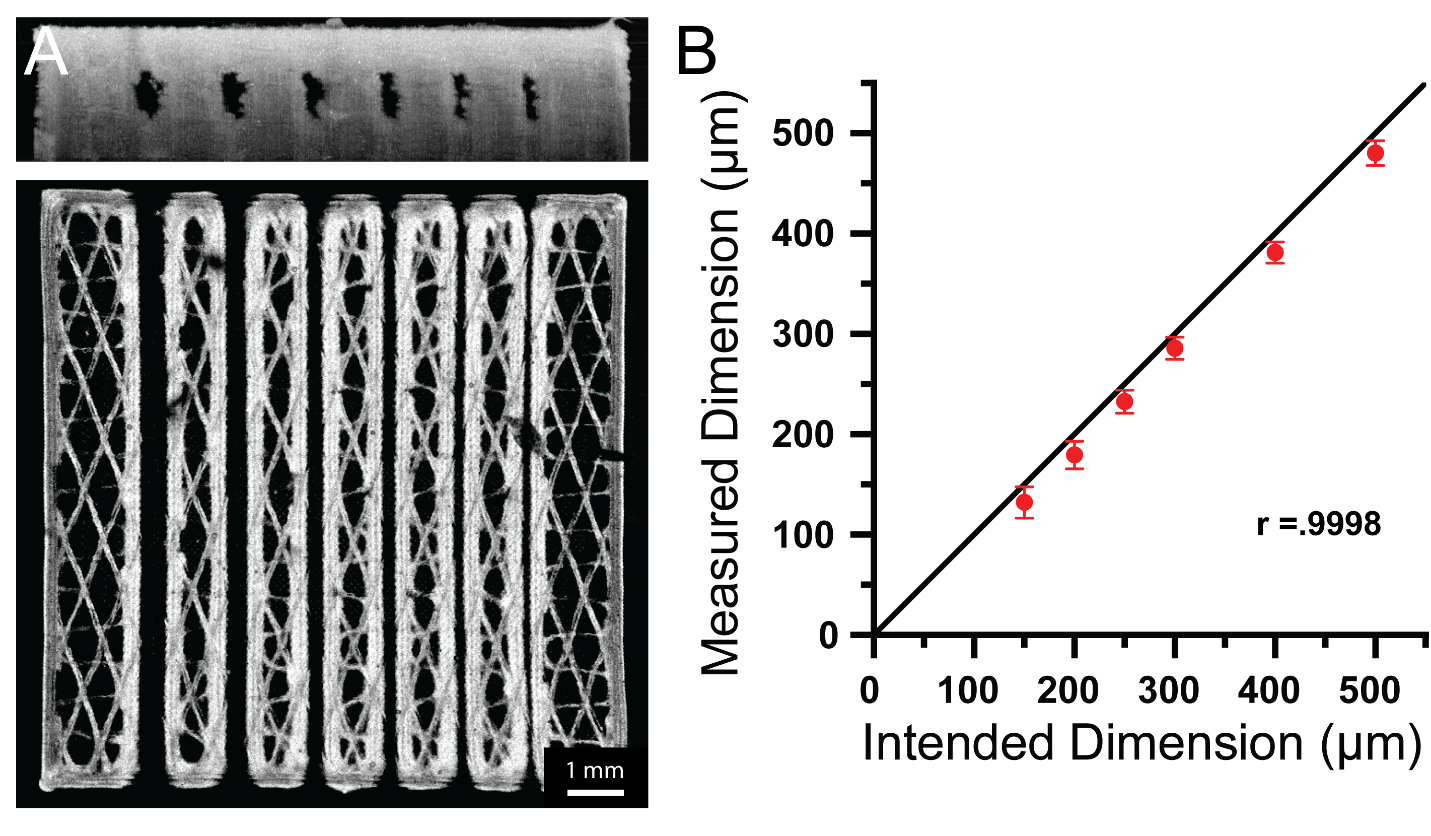
 **Figure S4| High contrast collagen print released from clear support retains its shape.** (**A**) An end on (above) and top down (below) view of a benchmark model printed using high contrast collagen I in clear support, after melting away the support. (**B**) Measured versus intended dimensions for the released print RMS error = 18.2884 µm (mean ± STD; n = 1 print with 9936 measurements, Pearson correlation coefficient = .9998).

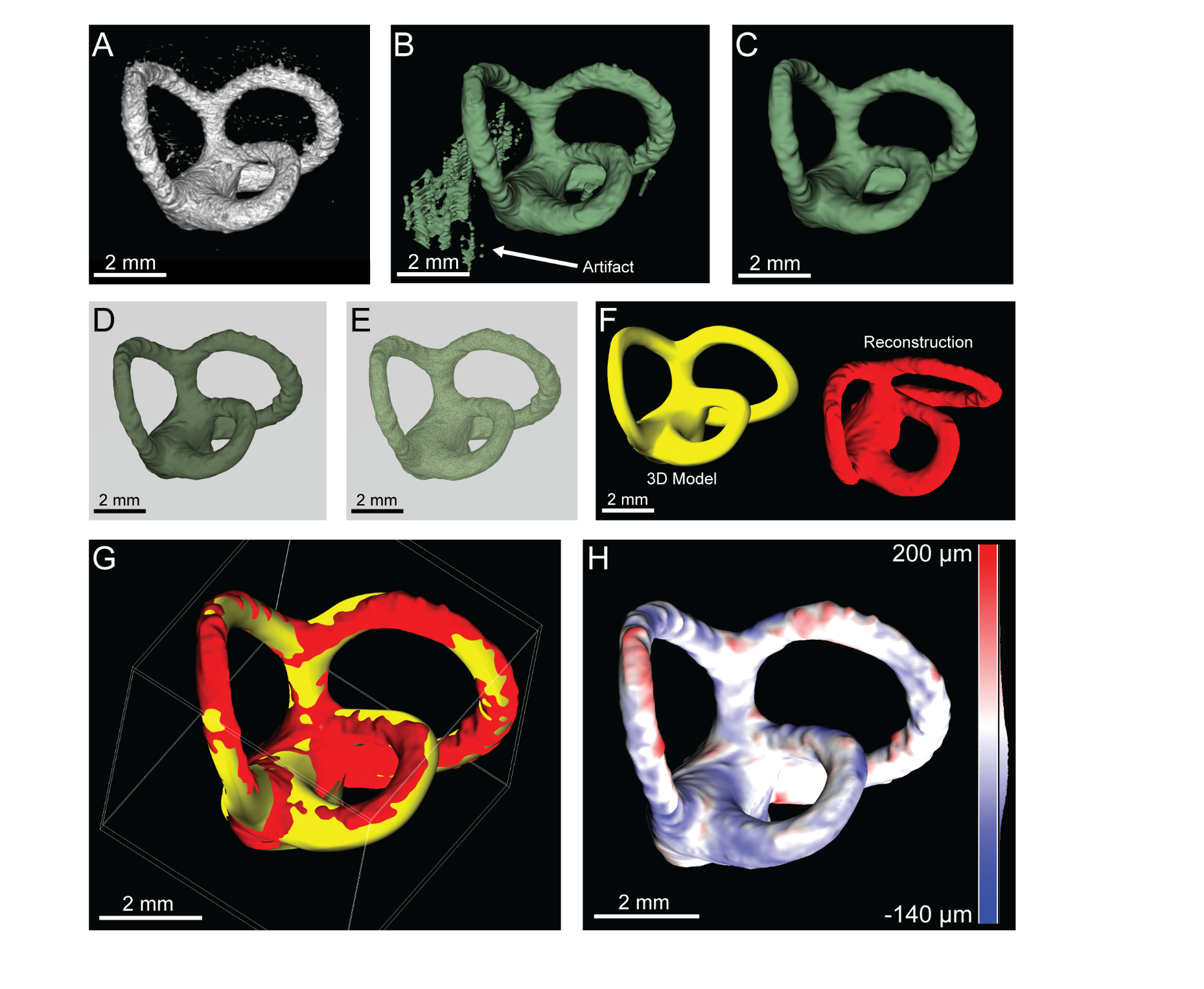
 **Figure S5| From composite OCT image to 3D reconstruction and gauging.** (**A**) A composite OCT image represented as a cloud of intensity data. (**B**) An initial segmentation of the OCT image data, with artifacts. (**C**) Segmentation corrected to remove artifacts, using raw OCT image data as reference. (**D**) Segmentation is exported as an STL with many polygons. (**E**) The STL mesh is decimated to decrease the number of polygons. (**F**) The original model (yellow) and the decimated reconstruction (red) are oriented incorrectly for comparison when loaded into CloudCompare. (**G**) After manual translation and rotation, the two models are automatically registered. (**H**) After registration, the two models can be compared to assess deviations from the original model (red = oversize, blue = undersize).

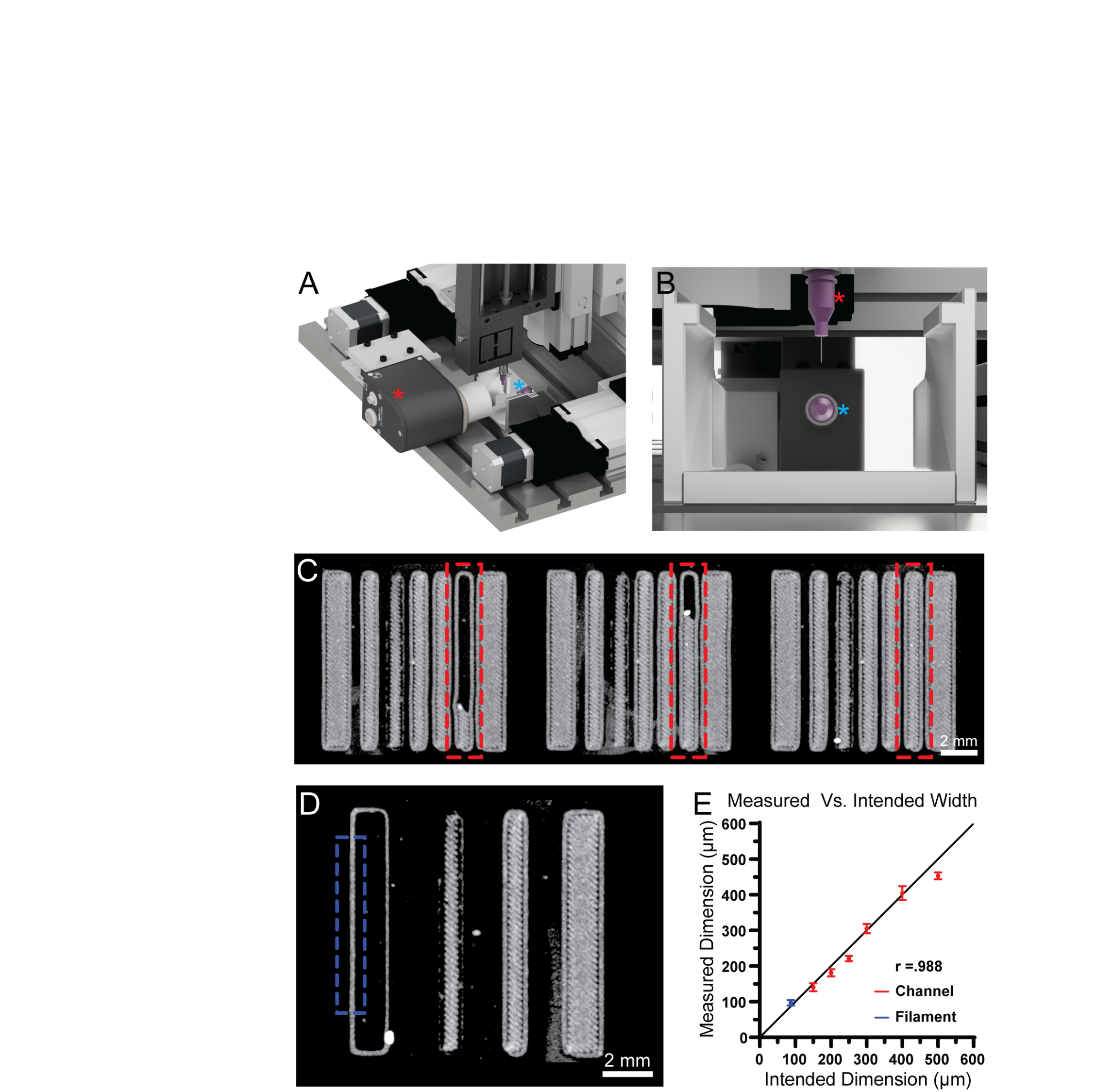

**Figure S6| Imaging setup for acquiring OCT during the printing process.** (**A**) An isometric view of the setup showing the OCT scanhead (red *) on its side with the objective aimed at a mirror mounted at a 45 degree angle (cyan *). (**B**) A view aligned with the OCT objective aimed at the mirror. The needle seen above (red *) can be viewed end on in the mirror (cyan *). (**C**) A time-lapse representation of OCT images acquired during printing. The region bounded by the red dashed rectangle is actively being printed. (**D**) Imaging with the OCT allows for measurements of features, such as walls and filaments, as they are being assembled. (**E**) Using the images acquired while printing measurements can be taken to verify the dimensions RMS error = 22.7909 µm (mean ± STD; n = 467 measurements of one channel/filament, Pearson correlation coefficient = 0.988).
